# Mechanisms of activation and desensitization of full-length glycine receptor in membranes

**DOI:** 10.1101/788695

**Authors:** Arvind Kumar, Sandip Basak, Shanlin Rao, Yvonne Gicheru, Megan L. Mayer, Mark S.P Sansom, Sudha Chakrapani

**Author notes:** Correspondence and requests for materials should be addressed to S.C.

## Abstract

Glycinergic synapses play a central role in motor control and pain processing in the central nervous system. Glycine receptors (GlyR) are key players in mediating fast inhibitory neurotransmission at these synapses. While previous high-resolution structural studies have provided insights into the molecular architecture of GlyR, several mechanistic questions pertaining to channel function are still unknown. Here, we present Cryo-EM structures of the full-length GlyR protein reconstituted into lipid nanodiscs that are captured in the unliganded (closed) and glycine-bound (open and desensitized) conformations. A comparison of the three states reveals global conformational changes underlying GlyR channel gating. The functional state assignments were validated by molecular dynamics simulations of the structures incorporated in a lipid bilayer. Observed permeation events are in agreement with the anion selectivity of the channel and the reported single-channel conductance of GlyR. These studies establish the structural basis for gating, selectivity, and single-channel conductance of GlyR in a physiological environment.

## Introduction

Glycine receptors (GlyRs), along with GABA_A_ receptors, are the principal determinants of fast inhibitory synaptic neurotransmission in the central nervous system (CNS)^1^. Glycinergic neurotransmission critically regulates motor coordination, sensory reflex activity, and respiratory rhythms. GlyR dysfunctions are associated with neuromotor deficiencies such as hyperekplexia and epilepsy, and underlie chronic inflammatory pain^2–4^. Agents that potentiate GlyR function produce analgesia making them attractive candidates for pain therapy^5^. GlyRs belong to the superfamily of pentameric ligand-gated ion channels (pLGIC) and assemble as either homopentamers of α subunits (α1−α4) or heteropentamers of α subunits in combination with β subunits (β1−β2). The GlyR has a conserved pLGIC architecture^6–10^ consisting of an extracellular domain (ECD), a transmembrane domain (TMD), and an intracellular domain (ICD). The ECD houses neurotransmitter binding pocket and the TMD governs the machinery for selective ion permeation. The ICD is important for receptor trafficking, synaptic clustering and is a central site for posttranslational modifications and regulation by endogenous and exogenous modulators^11–15^.

In the absence of glycine, an activating ligand for GlyR, the channel primarily resides in the resting (closed) conformation. Binding of glycine induces a global conformational change in the GlyR that leads to the opening of the channel pore lined by the M2 helices in the TMD. The open conformation is transient and the channel quickly transitions to a glycine-bound, desensitized conformation. Over the last few years, crystallographic and cryo-electron microscopy (cryo-EM) studies of GlyR have provided the first snapshots of the channel in distinct conformations^6,16^. These landmark studies revealed a conserved mechanism for channel activation involving a ligand-induced global twist in the ECD and TMD leading to channel opening. However, there are inconsistencies between these structures and previous work, particularly concerning charge selectivity, permeation, and single-channel conductance. One concern is the pore of the previously reported open GlyR conformation (PDB_ID: 3JAE) is considerably wider than what has been seen in other anionic pLGIC structures, particularly at the selectivity filter^6,17^. Molecular dynamic (MD) simulations with this structure predict a channel conductance that is higher than observed in electrophysiology studies and only partially blocked by the complete pore-blocking agonist picrotoxin^18^. These features are in contrast to electrophysiological measurements suggesting that the pore opening may have been exaggerated. Moreover, in MD simulations performed in the absence of backbone restraints, the open structure has been shown to rapidly collapse at the pore^19^. One possible cause of these discrepancies is that the previous structures were solved with a truncated ICD (consisting of 75-125 residues) to improve sample stability. The ICD, besides its role in trafficking, has also been implicated to alter single-channel conductance and gating^3,20^. Furthermore, the lipid environment is an essential component of membrane protein stability. Past structures have been determined in a detergent environment, though presently lipid nanodiscs are generally accepted to better mimic a natural lipid environment. It seems likely that both the ICD and lipid environment are important for maintaining the structural integrity of the channel during gating conformational cycles. We therefore determined high-resolution structures of the full-length GlyR in a membrane nanodisc environment by single-particle cryo-electron microscope (Cryo-EM). This allowed us to elucidate distinct physiological functional states of GlyR that give a complete mechanistic description of gating.

## Results and Discussion

### Biochemical characterization and structure determination

The zebrafish GlyRα1 exhibits robust macroscopic currents in response to application of glycine with an EC_50_ of 100 μM with current amplitude saturating beyond 1 mM^21^. In the continued presence of glycine, the currents desensitize within minutes. Upon application of picrotoxin (PTX), a convulsant alkaloid and a known open-channel blocker of most anionic pLGICs, GlyR currents are strongly inhibited with an IC_50_ of 5-9 μM, and a near complete inhibition is achieved at concentrations >1 mM^22^. The block is reversible, and previous studies have demonstrated that PTX block prevents the channel from desensitizing^23^. Our strategy to capture the GlyR in the resting, open, and desensitized conformations, was therefore to prepare samples of GlyR in the absence of a ligand, in the presence of both glycine and PTX, and in the presence of glycine, respectively.

The full-length glycineα1 receptor (GlyR) gene from Zebrafish was codon-optimized, cloned into pFastBac1 plasmid, and expressed in *Spodoptera frugiperda* (Sf9) cells (Supplemental Figure 1A). The sample stability was improved by incorporating a lipid mixture consisting of soybean polar extract (asolectin) and cholesterol hemisuccinate (CHS) during detergent solubilization and purification. The purified pentameric population of GlyR was then reconstituted into asolectin nanodiscs with the E3D1 membrane scaffolding protein. The nanodisc samples were used for single-particle cryo-EM analysis (Supplemental Figure 2). The samples were imaged in the absence of ligand (GlyR-Apo), in the presence of 5 mM glycine (GlyR-Gly), and in the presence of 5 mM glycine and 3 mM picrotoxin (GlyR-Gly/PTX). Three dimensional cryo-EM reconstructions with imposed C5 symmetry for the three sample sets led to density maps with nominal resolutions of 3.33, 3.55 and 3.63 Å, respectively (Supplemental Figure 3). In each case, the final reconstruction contained density for the entire ECD and TMD as well as a select region of the ICD, which were used for model building and refinement (Supplemental Figure 4). The model includes residues Pro31-Phe341, and Lys394-Gln444. The region between Phe341-Lys394, which is a part of the M3-M4 loop, is unstructured and appears as segmented density that is not conducive to model building. The density surrounding the TMD corresponds to the nanodisc belt comprising of the helical segments of the membrane scaffolding protein and lipid bilayer within the enclosure (Supplemental Figure 1C). There are three sets of additional non-protein densities; one appears as an extension from Asn62 corresponding to N-glycans and two are lipid densities in the vicinity of M4 (Supplemental Figure 2C). The glycan density at Asn62 was also observed in previous structures^6,16^ and corresponds to the single glycosylation site shown in GlyRα1. Mutations at this site have been reported to affect surface expression of functional GlyRs by preventing their exit from the endoplasmic reticulum^24^. The lipid at the extracellular end was modelled as phosphatidylcholine and the lipid at the cytosolic end was modelled as phosphatidylinositol-4,5-bisphosphate (PIP_2_) (Supplemental Figure 2C). The PIP2 density is in a similar location to that observed in a recent GABA_A_R structure^8^.

The overall architecture, consistent with the previous GlyR structures and other pLGIC structures^6–10^, consists of a symmetrical arrangement of the five subunits with dimensions of 125 Å in height and 80 Å in diameter. Each subunit consists of a twisted β-sheet of ten strands forming the ECD, four α-helical strands comprising the TMD, and a primarily unstructured region between the third and fourth TM helices forming the ICD (Supplemental Figure 1B). The TM helices splay outward toward the extracellular end of the bilayer. The M4 helix extends out of the putative bilayer boundary (referred to as post-M4), by 3 turns of the helix and makes contact with the ECD. The ICD is the least conserved region among the pLGIC family and varies both in length as well and amino acid sequence. In particular, it is expected that there is extensive structural variation between anionic and cationic pLGIC, in line with the distinct roles played by this domain in the two channel classes. In GlyR, the M3-M4 linker extends as a continuous stretch of α-helix from the C-terminal end of M3 up to four turns (referred as post-M3). Beyond this, it is predicted to be an unstructured region that eventually terminates with a short α-helix (referred as pre-M4) that extends into the M4 heix. In contrast, the ICD in cationic pLGICs consists of an unstructured loop (post-M3 loop), followed by an amphipathic MX helix that runs parallel to the membrane, an unstructured region that ends in a long α-helix MA which appears as a single continuous helix with M4^25,26^. In the resting conformation, both the post-M3 loop and the MA helix occlude the ion permeation pathway, and ligand-binding elicits large conformational changes in this region to create a lateral portal for ion exit^9,27^. In the absence of extensive structured motifs, we speculate that the ICD in anionic channels may not impose such physical barriers to permeation.

### The permeation pathway

The GlyR-Apo, GlyR-Gly/PTX, and GlyR-Gly structures reveal distinct conformational states of the ion permeation pathway, which originates at the ECD, extends through the TMD, and terminates shortly within the ICD (Fig 1A). Within the TMD, the pore is lined by the M2 helix from each subunit in an arrangement that appears more cylindrical in GlyR-Apo and becomes progressively funnel shaped, tapering toward the intracellular end in GlyR-Gly/PTX and GlyR-Gly. In the GlyR-Apo, the pore is relatively narrow and constricted prominently at Leu9′ (pore radius ∼ 1.4 Å) and Thr13′ (pore radius ∼ 2.2 Å). The pore dimensions at positions Ala20′ and Pro-2′ are also below the Born radius for the solvated chloride ion which is 2.26 Å (the Pauling radius for chloride ion is 1.81 Å)^28^ and are therefore likely barriers to ion permeation (Fig 1B). The M2 helices show partial unwinding between Gly17′ and Ala20′, a feature that is consistent with the dynamic behavior of this region previously noted in NMR studies of the isolated GlyR TMD^29^. In the GlyR-Gly/PTX structure, the M2 helices are oriented outward with the Leu9′ rotated away from the central axis (Fig 1C). The pore radius profile (created with the PTX ligand removed from the pore) shows an expanded ion permeation pathway at each of the constriction points seen in GlyR-Apo. In GlyR-Gly, the pore is similar in radii to that of the GlyR-Gly/PTX but reveals a notable constriction (pore radius ∼ 1.8 Å) at the level of Pro-2′ due to slight bending of the intracellular end of M2. The Pro-2′ position is part of the charge selectivity filter in anionic channels comprising of a conserved stretch (Pro-2′, Ala-1′, and Arg0′)^30,31^ and mutations are associated with hyperekplexia^32,33^. While Pro-2′ and Ala-1′ face the lumen of the pore, Arg0′ faces away. Previous studies have shown that both charge as well as the side-chain conformation of Arg0′ are determinants of anion selectivity^34^.

**Figure 1.**
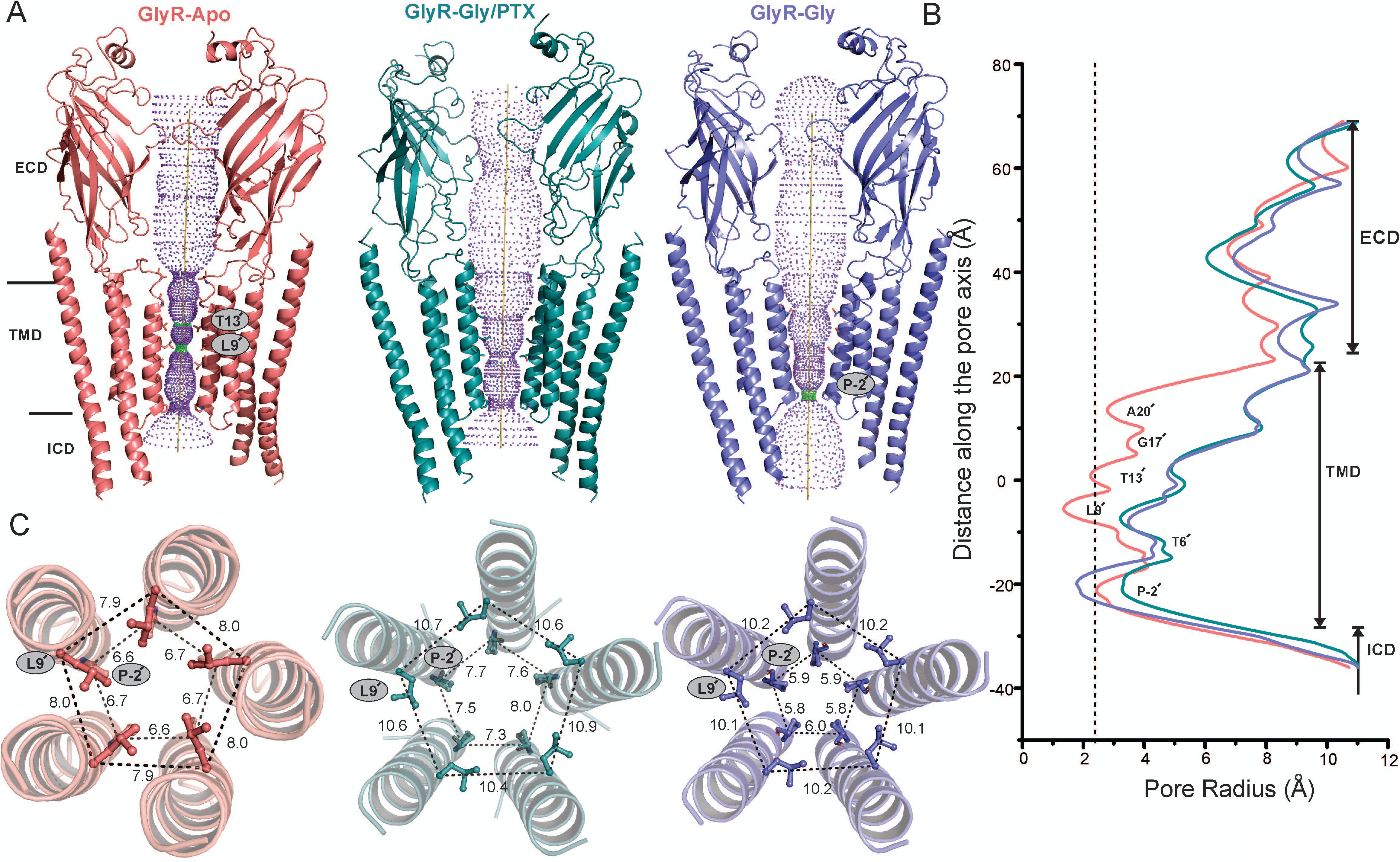
Cryo-EM structures of full-length GlyR in multiple conformational states. A) Ion permeation pathway generated with HOLE^56^ for GlyR-Apo (salmon red), GlyR-Gly/PTX (deep teal) and GlyR-Gly (slate blue). For clarity, the cartoon representation of only two non-adjacent subunits are shown. Green and purple spheres define radii of 1.8-3.3 Å and > 3.3 Å, respectively. The residues located at various pore constrictions are shown as sticks. B) The pore radius is plotted as a function of distance along the pore axis. The dotted line indicates the approximate radius of a hydrated chloride ion, which is estimated at 2.26 Å. C) A view of M2 helices from the extracellular end for the three GlyR conformations. Positions Leu9′ and Pro−2′ are shown in ball-and-stick representation and the corresponding distances between Cα are given in Å.

In addition to the physical dimensions of the pore, the hydrophobicity of the pore-lining residues plays a key role in dictating the conductance state of the channel. In some cases, a pore that is not narrow enough for steric occlusion may still impede ion permeation due to the hydrophobicity of the pore-lining residues that may disfavor water molecules, causing local dewetting that leads to a higher energetic barrier for water and ion permeation^35^. We assessed the combined effect of both the pore radii and hydrophobicity on pore dewetting and presence of hydrophobic gates using a previously developed simulation-free heuristic model that was derived using machine learning from MD simulations of nearly 200 ion channel structures^36^. A pore surface profile based on the radii of the permeation pathway and hydrophobicity of the pore-lining amino acid side chains was estimated by CHAP^37^ (Fig 2A). On the plot of the local pore radii at each pore-lining residue against its side-chain hydrophobicity, the dotted line marks the divide of the hydrophobicity-radii landscape into regions of high and low likelihood of pore wetting (Fig 2B). The sum of shortest distances (Σd) from positions falling in the low-likelihood region to the line is used as a heuristic score which allows a prediction for the likelihood of the conformation corresponding to a dewetted and non-conductive state. A cut-off value of Σd> 0.55 is used for predicting a non-conductive conformation. In the GlyR-Apo, several residues (Pro-2′, Ile5′, Leu9′, and Thr13′) fall below the line, leading to an overall heuristic score of Σd = 1.61. In the GlyR-Gly/PTX, there are no residues below the line and the score is 0.07. On the other hand, in the GlyR-Gly structure, Pro-2′ is well below the line with Val1′ lying close and the score is 0.80. These findings suggest that the pore in Gly-Gly/PTX structure is likely to be open, and those in the GlyR-Apo and GlyR-Gly are closed. The electrostatic map shows the positively charged cluster at the intracellular end of M2 and the regions of the ICD close to the TMD interface (Fig 2C). The Arg0′ that is implicated to be the selectivity filter in anionic pLGIC lies on the non-pore facing side of M2 at the interface of the TMD and ICD.

**Figure 2.**
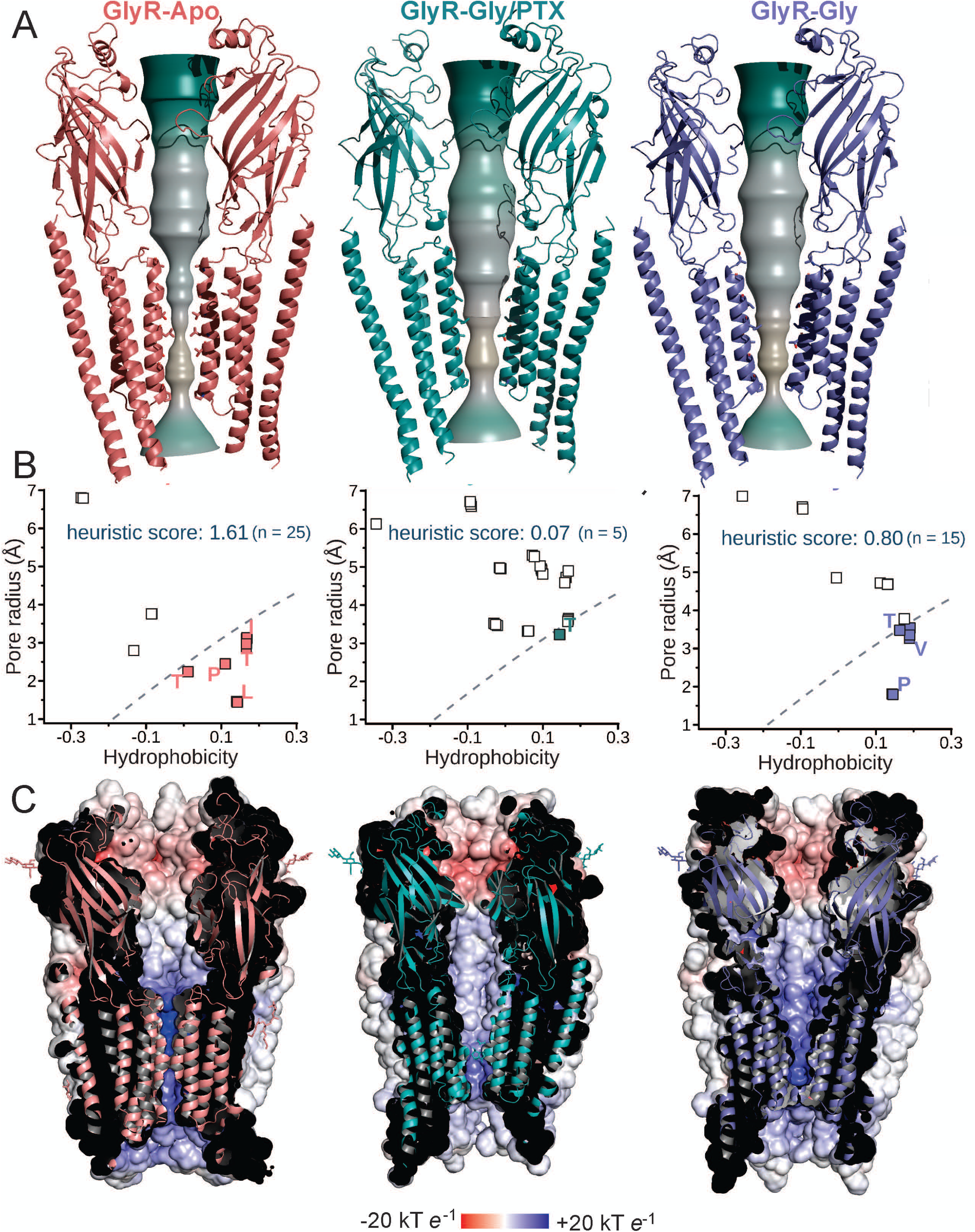
Assessment of pore properties. A) Pore surface through each channel structure, colored by hydrophobicity (from hydrophilic-green, to hydrophobic-yellow) as estimated by CHAP^3^ based on the experimental hydrophobicity values of pore-lining amino acid side-chains. B) Assessment of the likelihood of pore closure by an energetic barrier corresponding to dewetting at a hydrophobic region, evaluated according to a heuristic method based on simulation of water behavior in ∼200 ion channel structures^36^. Any identified pore-lining sidechains are shown as points and were mapped onto the prediction grid with their local pore hydrophobicity and radius as coordinates. The subset of those falling below the dashed classification line were used to calculate a heuristic score. A cut-off of > 0.55 had been used to predict that a channel structure would contain a hydrophobic barrier to water and ion permeation. C) The vertical slice-through of the receptor showing the electrostatic surface potential generated using the APBS tool plug-in in PyMOL^57^ along the ion-conducting pathway for GlyR-Apo, GlyR-Gly/PTX and GlyR-Gly conformations.

### Ligand-binding sites

The GlyR-Gly and GlyR-Gly/PTX structures were solved in the presence of 5 mM glycine (Fig 3A) and the 3D reconstructions reveal an additional small density in the canonical neurotransmitter binding pocket wedged at the subunit interface in the ECD (Fig 3B). An unambiguous assignment of a small ligand such as glycine is not possible at this resolution, however, since the density was not seen in GlyR-Apo, and the location matched with the glycine density in the α3-GlyR crystal structure^38^, we modelled the glycine ligand accordingly. The neurotransmitter binding pocket is lined by several conserved aromatic residues and bulky side-chains, from both the principal (Phe123, Phe 183, Tyr226, Thr228, and Phe231) and complementary subunits (Phe68, Arg89, Leu141, Ser145, and Leu151). Mutational studies have shown that a significant impact on agonist affinity ensues upon perturbation of this pocket^39,40^. An overlay of GlyR-Apo and GlyR-Gly shows substantial conformational changes that lead to a rearrangement of the residues in the binding pocket (Fig 3C). The major motion includes an inward movement of Loop C (carrying Tyr226, Thr228, and Phe231) and Loop B (Phe 183). The tip of Loop C (measured at Cα of Thr228) is displaced by 6.2 Å towards the neurotransmitter binding pocket thereby shrinking the pocket and potentially occluding water molecules. The inward or “capping” motion of Loop C is well-described in pLGIC literature and is associated with agonist-binding^41^ and is also observed in the recent structures of the unliganded and agonist-bound conformations^6,9,17,27,42^.

**Figure 3.**
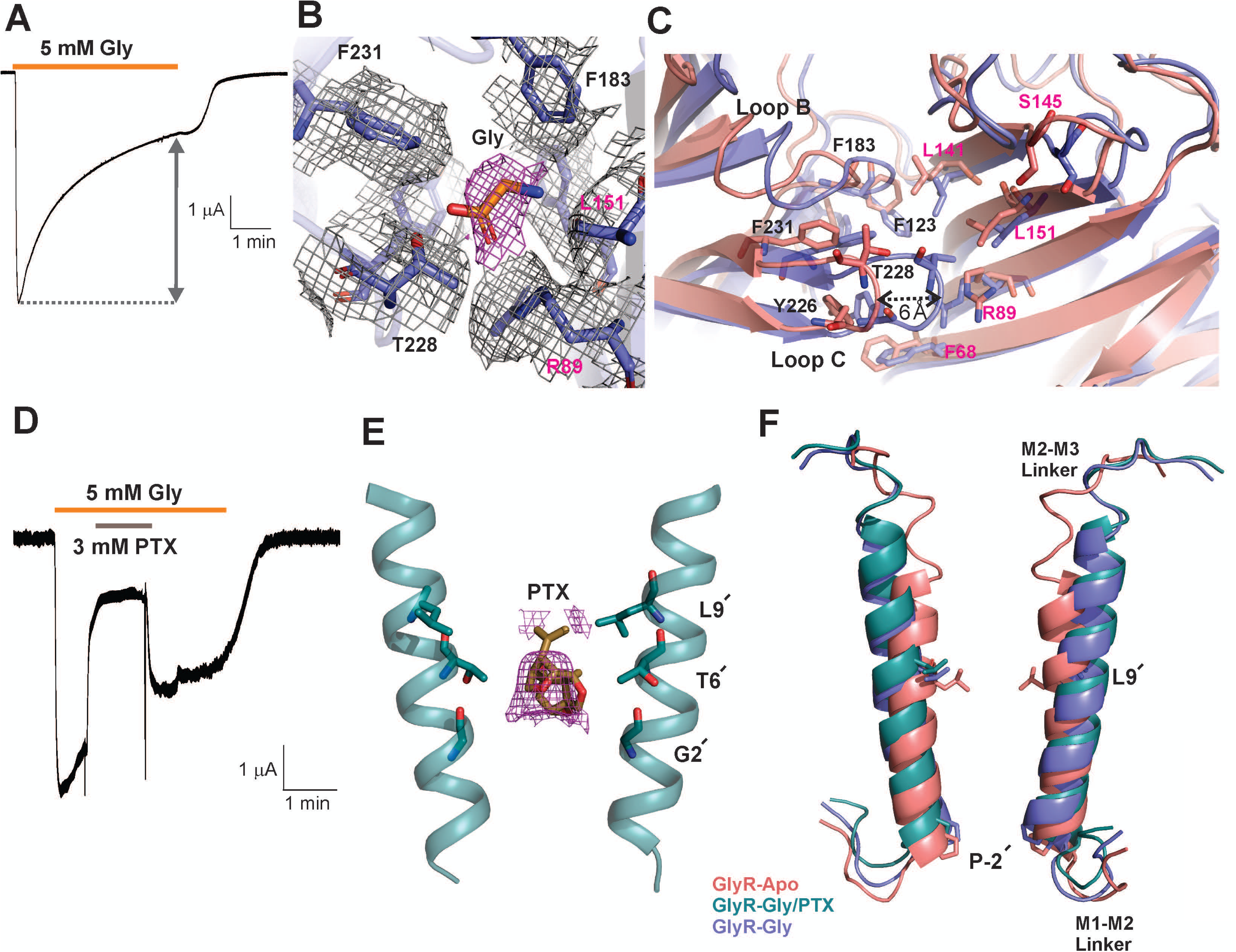
Conformational changes underlying GlyR gating. A) Two-electrode voltage clamp (TEVC) recording of GlyR expressed in *Xenopus* oocytes. Currents were elicited in response to application of 5 mM Glycine with a holding membrane potential of −60 mV. Robust inward currents are observed that desensitize in the presence of glycine. B) Cryo-EM density segments for neurotransmitter binding site residues (gray) and glycine ligand (pink) as seen in the GlyR-Gly structure. C) Comparison of the neurotransmitter binding site for the GlyR-Apo and GlyR-Gly conformations. The residues that are involved in neurotransmitter binding are shown in sticks. D) A TEVC recording showing the effect of 3 mM PTX block on GlyR current evoked by application of 5 mM glycine with a holding membrane potential of −60 mV. Upon wash-off of PTX, the amplitude of recovered currents reflects the population of channels still conductive. E) Cryo-EM map showing the density for PTX (pink) bound in the pore of GlyR-Gly/PTX. The interacting residues are shown in stick representation. For clarity, only parts of M2 for two diagonal subunits are shown. F) A close-up of the M2 conformations is shown upon aligning the three GlyR conformations. Positions Leu9′ and Pro-2′ are shown as sticks.

In the presence of glycine, PTX reversibly blocks GlyR currents (Fig 3D). In the GlyR-Gly/PTX reconstruction, in addition to the glycine density at the neurotransmitter binding pocket, an additional density was seen in the pore at the mid-level of M2 nestled between Gly2′ and Leu9′ (Fig 3E). This density was not observed in the other two conditions and likely corresponds to a PTX molecule. A recently solved structure of GABA_A_R also contains a PTX molecule at the same location and was used to guide our precise orientation of the molecule.^38^ The hydrophobic isoprenyl end of PTX is oriented toward the Leu9′ and the hydrophilic end closer to the Thr6′. Mutational studies have shown that perturbations at these positions affect PTX binding in anionic pLGICs^43–45^. An inspection of the M2 helices upon global alignment of all three conformations show that the binding site for PTX does not physically overlap with the region undergoing constriction in the GlyR-Gly (Pro-2′). Therefore, it appears that PTX exerts an allosteric effect on the intracellular gate Pro-2′ and prevents it from closing in the presence of glycine.

### Glycine induced global conformational change

When compared to GlyR-Apo, the glycine-bound structures undergo an anti-clockwise twist of the ECD around the pore axis (when viewed from the extracellular end). The ECD twist is accompanied by an inward motion of Loop C described above (Fig. 4A). In addition, there is major repositioning of the interfacial loops, particularly the β1-β2 loop and Cys-loop (β6-β7) from the principal subunit and Loop F (β8-β9) on the complementary subunit. The TMD helices are rotated clockwise and tilted outward in the GlyR-Gly/PTX and GlyR-Gly structures compared with GlyR-Apo, leading to an iris-like opening of the TMD (Fig. 4B). In addition to the M2 helices moving outward to open the pore, the M2-M3 linker is also pulled away from the pore axis. This twisting motion is similar to the previously observed changes in GlyR and other pLGICs^6,46^. The described motions have a substantial effect on interactions between subunits at both the ECD and TMD interfaces as well as intra-subunit interactions between the ECD and TMD (Fig 4C). In GlyR-Apo, the Cys-Loop, β1-β2 linker, and Loop F from the ECD are in intimate contact with the M2-M3 linker, pre-M1, and post-M4. In addition, there are several interactions between M1, M3 and M4 helices between two adjacent subunits. In the glycine-bound conformations, these interactions are substantially reduced due to the ECD rotation and the TMD expansion at the extracellular end. The overall buried surface area in GlyR-Apo is 34, 500 Å^2^ and reduces to 26, 830 Å^2^ in GlyR-Gly/PTX Å^2^, and 26, 130 Å^2^ in the GlyR-Gly. The interaction network at the domain interfaces are crucial signal transduction elements and, unsurprisingly^47,48^, mutational perturbations are reported to significantly alter function, and in a number of cases, related to disease pathology^2,49^.

**Figure 4.**
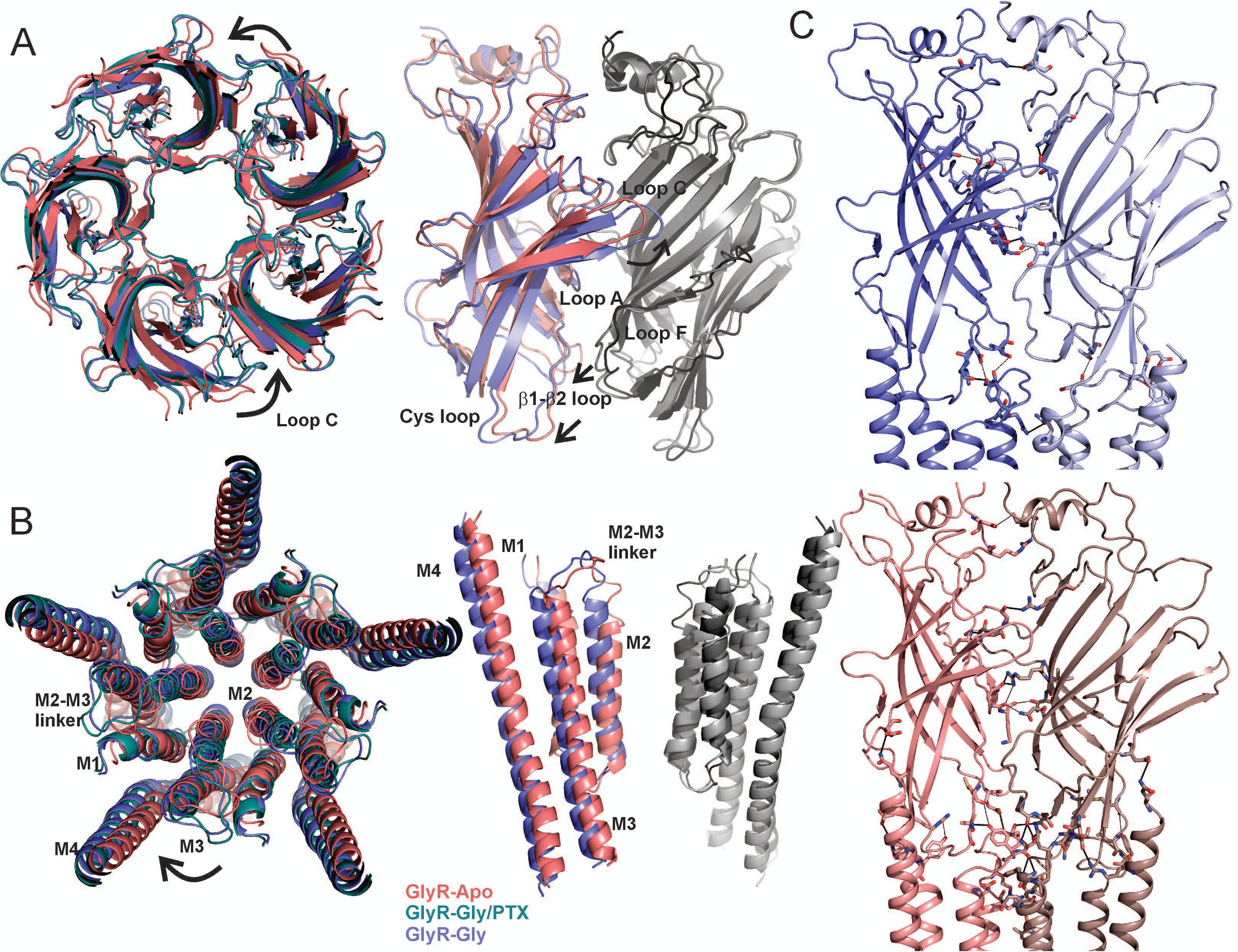
Global conformational changes and altered inter-domain interactions during GlyR gating. A) A view of the ECD from the extracellular end upon aligning GlyR-Apo, GlyR-Gly/PTX, and GlyR-Gly (*left*). Glycine induced anti-clockwise rotation of the ECD is highlighted by black arrows in the vicinity of Loop C. A side-view of the ECD interface in GlyR-Apo and GlyR-Gly. The principal subunits are colored (GlyR-Apo: salmon red and GlyR-Gly: slate blue) while the complementary subunits are shown in shades of gray. B) A view of the TMD from the extracellular end upon aligning GlyR-Apo, GlyR-Gly/PTX, and GlyR-Gly (*left*). There is a clockwise rotation of the TMD helices with an outward expansion resembling the opening of an iris. A side-view of the TMD in GlyR-Apo and GlyR-Gly conformations C) Interactions at the subunit and domain interfaces between ECD-ECD, ECD-TMD, and TMD-TMD. The residues involved in the interactions are shown in sticks and interactions are highlighted by black dashes.

### The M4 conformation and the effect on intra-subunit and inter-subunit cavities

Glycine-induced conformational changes within the TMD leads to expansion of the TM helices. In GlyR-Apo, M4 helices are oriented closer to the rest of the TM helices within the subunit. The post-M4 region extends three turns above M4 and extends above the putative membrane and interacts with the pre-M1 region and the β8-β9 strand of the ECD (Fig 5A).These regions are implicated in relaying signal from the ECD to the TMD. In the GlyR-Gly the residues involved in these interactions are placed farther away. At the intracellular end, the M4-pre-M4 helices (intracellular extension) are bent in the vicinity of Pro419 and positioned such that there is a favorable interaction with the lipid molecule (modeled as PIP2). In the GlyR-Gly structure, the M4 and pre-M4 helices are further straightened, and there is no obvious lipid density at this position, implying a potential loss of interaction (Fig 5B). The movement of post-M3 helices are in the same direction as the pre-M4, and is potentially coupled by the unstructured loop that is absent from our structures. Among other related movements at the intracellular end, is the movement of the M1-M2 linker (Fig 5B). In the GlyR-Gly structure, the M1-M2 linker is moved away from the pore axis as a consequence of (or to facilitate) the pinching of M2 helices at Pro-2′. In agreement, both the length and the sequence of the M1-M2 linker affect GlyR gating and desensitization, presumably by affecting the positioning of the selectivity filter region (Pro-2′, Ala-1′, and Arg0′)^50,51^. Another significant consequence of structural changes at the TMD and ICD are the appearance of new intra-subunit and inter-subunit cavities at both the extracellular and intracellular ends of the TMD (Fig. 5C). GlyRs are targets to numerous endogenous and exogenous ligands that target TMD cavities. A change in the volume and polarity of these cavities implicates a role in state-dependent effects of the modulators on channel function^52,53^. Also of note, is the role of the ICD in imparting sensitivity to allosteric ligands in GlyR modulation^54^.

**Figure 5.**
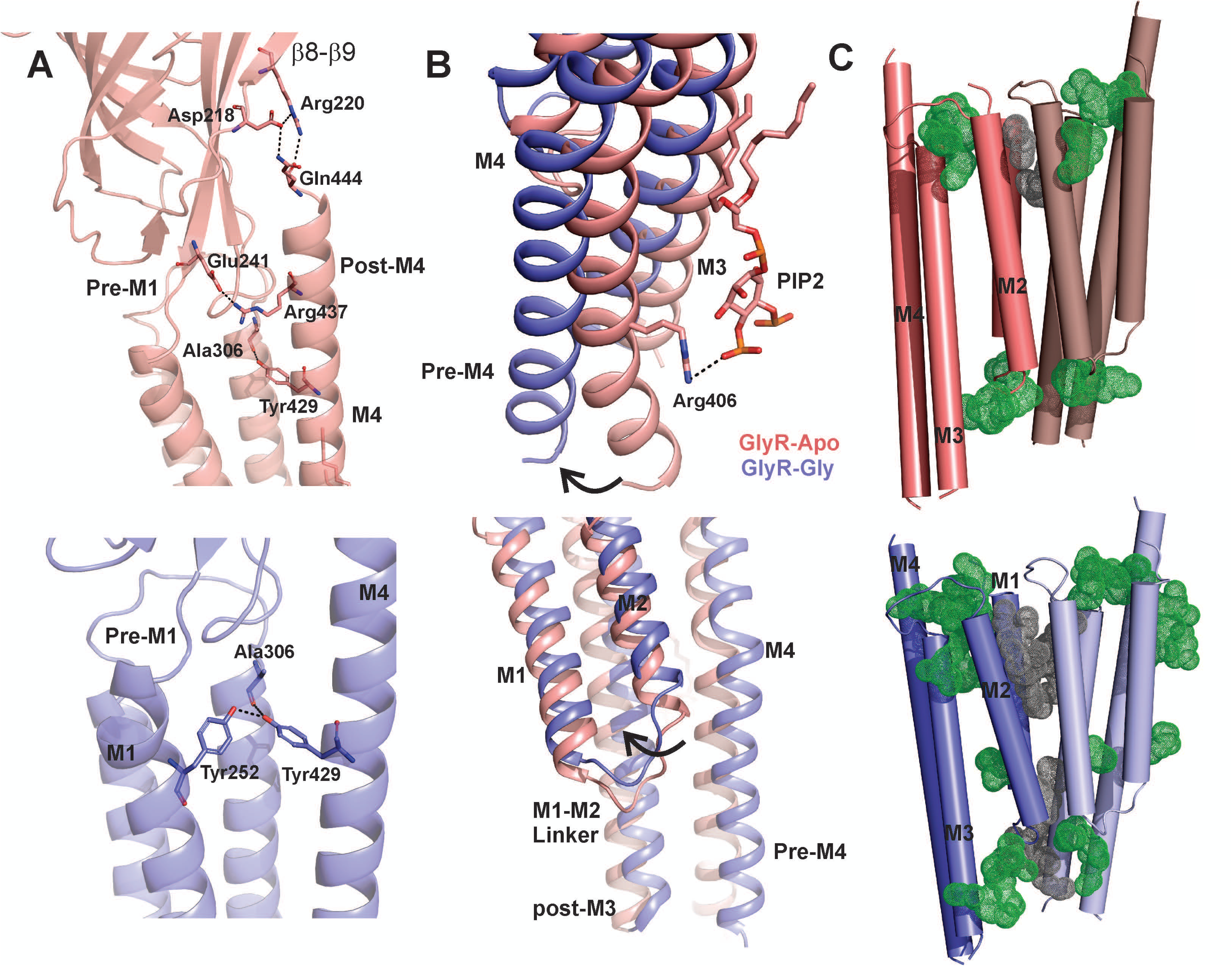
Conformational changes in the M4 helix and the effects on internal cavities. A) Interaction of the post-M4 region with the ECD in GlyR-Apo (*top*) and GlyR-Gly (*bottom*) conformations. The residues involved in the interactions are shown in sticks. B) An overlay of GlyR-Apo and GlyR-Gly reveals the extent of conformational change in the ICD formed by the pre-M4 region (*top*) and the M1-M2 linker and the post-M3 region (*bottom*). The direction of movement is highlighted by black arrows. A modeled PIP2 lipid molecule is in a potential interaction with Arg406. There was no clear density seen for the lipid in the GlyR-Gly reconstruction. C) A comparison of the inter-subunit (gray dots) and intra-subunit (green dots) cavities in GlyR-Apo (*top*) and GlyR-Gly (*bottom*) predicted using Fpocket algorithm^58^. The cavities predicted on the surface were removed for clarity.

We compared the full-length GlyR states with the previously solved GlyR cryo-EM conformations by aligning GlyR-Apo with StryR (strychnine-bound), GlyR-Gly/PTX with Gly (glycine bound-open), and GlyR-Gly with IVM (glycine/invermectin bound)^6^. While the structures overlap well (Supplemental Figure 5), the most notable differences for each alignment pair are in the TMD and the interfacial interactions. The buried surface area of StryR is 27, 392 Å^2^ with greater numbers of inter-domain interactions compared with that of 34, 500 Å^2^ in GlyR-Apo. The structural elements comprising of the ICD, namely the M1-M2 linker and the post-M3 and pre-M4 helices adopt distinct conformations. Additionally, in comparison with the GlyR-Gly/PTX, the glycine-bound open state is much wider, particularly at the intracellular end where the charge selectivity filter of the channel resides (Arg0′). We believe that some of these differences may arise from truncation and the presence of a detergent environment.

### Assessment of conductance states by molecular dynamics simulations

By evaluating the hydrophobicity and radius of the transmembrane pore, the constrictions at Leu9′ and Pro-2′ were predicted to form energetic barriers to varying extents in each of the GlyR conformational states. To assess the behavior of water molecules and ions for the three conformations in a membrane environment, molecular dynamics simulations were carried out upon embedding the GlyR structures within a 1-palmitoyl-2-oleoyl-*sn*-glycero-3-phosphocholine lipid bilayer (Fig. 6). The picrotoxin molecule in the pore of GlyR-Gly/PTX structure was removed prior to equilibration in the membrane. Positional restraints were placed on the protein backbone during these simulations, preserving the overall experimentally-determined conformational state whilst permitting rotameric flexibility in amino acid side chains. Water molecules (using the TIP4P/2005 model) and ∼150 mM NaCl were included on either side of the bilayer. During simulations, the side-chain movements of the pore-lining residues caused fluctuations in the pore-radius but there were no major changes to the pore profile (Fig 6A). An analysis of simulated water density along the pore axis suggests that in GlyR-Apo is the pore is dewetted (*i.e.* devoid of water molecules) at Leu9′ (at ∼ 0 Å, conferring closure to water with an energetic barrier of ∼4 kJ mol^-1^. This barrier disappears in both the GlyR-Gly/PTX and GlyR-Gly structures. However, while the pore is hydrated in GlyR-Gly/PTX, a small energetic peak (∼2.5 kJ mol^-1^) appears at the Pro-2′ position (∼-20 Å) which corresponds to a maximum free energy value of ca. 1 RT (i.e. comparable to that of thermal fluctuations). While water permeation across the pore is essential for ion permeation, it is not necessarily sufficient. To assess ion conductance for the three states, in a second set of simulations, a transmembrane potential of −200 mV (*i.e.* negative at the cytoplasmic side) was applied at 500 mM NaCl concentration. As expected, no ion permeation events were observed for GlyR-Apo. Interestingly, while water permeates through GlyR-Gly/PTX, no chloride ions passed through the channel during 1 μs of total simulation time. By contrast, an average of ∼12 chloride permeation events were observed per 200 ns simulation trajectory of the GlyR-Gly/PTX structure, corresponding to an estimated single-channel conductance of ∼50 pS for chloride efflux, comparable in magnitude to measurements from single-channel experiments which showed a conductance of 80-88 pS^55^. We believe that some of the differences in the conductance estimate may be due to the missing residues in the ICD that may exert an effect on the single channel conductance through electrostatic mechanisms. Choride ions were able to pass through the Pro-2′ position even when the pore radius was just above 2 Å during our simulations, in which instances they transiently became partially dehydrated such that interactions with Pro-2′ from the five subunits were not all water-mediated. It is noteworthy that sodium ions do not pass past the selectivity filter region comprising of Arg0′(∼-20 Å), confirming the anionic selectivity for GlyR. Our computational assessment of the three GlyR conformations thus indicate that the GlyR-Gly/PTX structure should be assigned to an open state, whereas the GlyR-Apo and GlyR-Gly structures are non-conductive.

**Figure 6.**
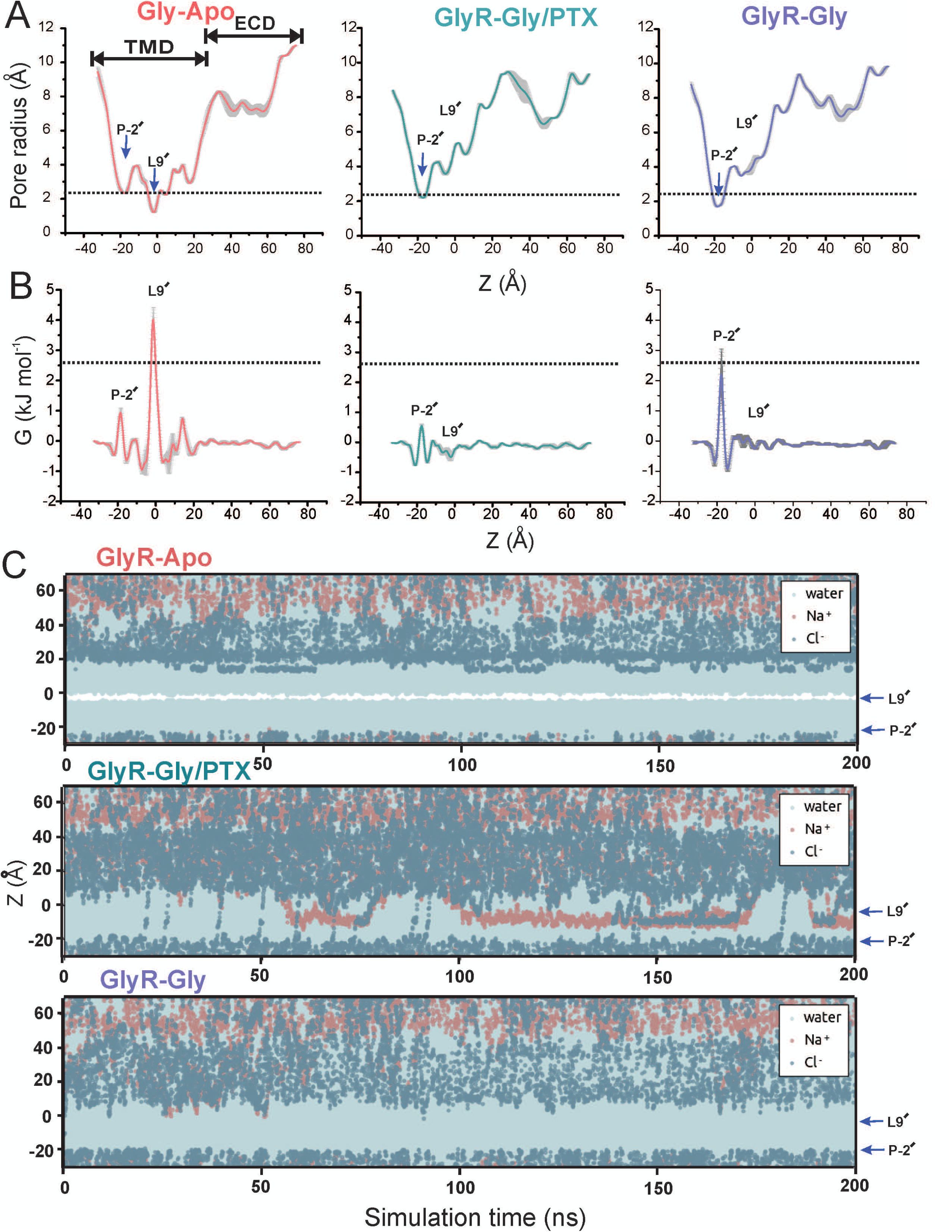
Molecular dynamics simulations of GlyR conformations. A) Mean pore radius profiles and standard deviations averaged across three independent 30 ns equilibrium simulations for GlyR-Apo (*left*), GlyR-Gly/PTX (*middle*), and GlyR-Gly (*right*) along the central pore axis. The final 20 ns of each 30 ns simulation trajectory was used to evaluate these profiles. The one-standard-deviation range between calculations (n = 3) are shown as a gray band. Major constriction sites are indicated and the dotted line denotes the radius of hydrated chloride ion. B) Corresponding mean water free energy profiles and standard deviations. Peaks in free energy profiles are highlighted. C) Trajectories along the pore (z) axis of water molecules and chloride ion coordinates within 5 Å of the channel axis inside the pore, in the presence of a −200 mV transmembrane potential difference (i.e. with the cytoplasmic side having a negative potential). One of five independent 200 ns replicates is shown for each structure. During these and the preceding simulations, positional restraints were placed on the protein backbone, in order to preserve the experimental conformational state whilst permitting rotameric flexibility in amino acid side chains. The energetic barriers due to the ring of Leu9′ and Pro-2′ are at z ∼ 0 Å and −20 Å, respectively.

Several previous simulation studies have noted the instability of the pLGIC conformations in the absence of protein backbone restraints. To assess the GlyR states, a separate set of 200 ns simulations were performed in which the restraints on the backbone were removed (Supplemental Figures 6 and 7). Under these conditions, while the fluctuations in the pore radii are understandably larger, the defining pore features of the three conformations were stably maintained, namely, the constrictions and energetic barriers to water at Leu9′ and Pro-2′ in GlyR-Apo and at Pro-2′ in GlyR-Gly. We found that the Leu9′ gate remains stably open in both of the GlyR-Gly and GlyR-Gly/PTX structures. Quite remarkably, in the simulations of GlyR-Gly/PTX the pore radius at Pro-2′ reduced toward that of the GlyR-Gly structure, leading to a constriction that would prevent ion permeation at this location. This further confirms that the open conformation of GlyR is transient and is trapped in this state due to the blocker which acts as a “foot-in-the-door” preventing the collapse of the intracellular pore.

## Conclusion

Here we present the first set of structures revealing the complete range of ligand-induced conformational changes in the full-length GlyR reconstituted in a lipid nanodisc environment. At the heart of pLGIC gating mechanisms is the signal transduction machinery that communicates ligand-binding events to the channel pore leading to channel opening. We show that glycine induces a global conformational change that encompasses the ECD, TMD, and the structured regions of the ICD. In the resting conformation, the GlyR has tightly coupled subunit and domain interfaces that weaken in response to glycine binding, leading to a relaxed open and desensitized conformations. Molecular dynamics simulations were used to assess the energy landscape of pore hydration and ion permeation which allowed us to annotate these structures as resting, open, and desensitized conformations. The single-channel conductance and ion selectivity predicted for the open conformation are in agreement with the wealth of previous functional studies. Overall, these studies set the stage for a detailed characterization of the functional modulation of this clinically important class of channels.

## Methods

### Electrophysiological recordings by two-electrode voltage-clamp (TEVC) in oocytes

The gene encoding zebrafish GlyRα1 (purchased from GenScript) was inserted into *pTLN* vector for expression in *Xenopus laevis*. To linearize the DNA, the plasmid was incubated with *Mlu1* restriction enzyme at 37 °C overnight. The mRNA was synthesized from linearized DNA using the mMessage mMachine kit (Ambion) per instructions in the manufacturer’s manual. The RNA was then purified with RNAeasy kit (Qiagen). About 2–10 ng of mRNA was injected into *X. laevis* oocytes (stages V–VI) and experiments were performed 2-3 days after injection. For control experiments to verify that no endogenous currents were present, oocytes were injected with the same volume of water. Dr. W. F. Boron kindly provided oocytes used in this study. Female *X. laevis* were purchased from Nasco. Animal experimental procedures were approved by Institutional Animal Care and Use Committee (IACUC) of Case Western Reserve University. Oocytes were maintained at 18 °C in OR3 medium (GIBCO-BRL Leibovitz medium containing glutamate, and 500 units each of penicillin and streptomycin, with pH adjusted to 7.5 and osmolarity to 197 mOsm). TEVC experiments were performed on a Warner Instruments Oocyte Clamp OC-725. Currents were sampled and digitized at 500 Hz with a Digidata 1332A, and analyzed by Clampfit 10.2 (Molecular Devices). Oocytes were clamped at a holding potential of −60 mV and solutions were changed using a syringe pump perfusion system flowing at a rate of 6 ml/min. The electrophysiological solutions consisted of (in mM) 96 NaCl, 2 KCl, 1.8 CaCl_2_, 1 MgCl_2_, and 5 HEPES (pH 7.4, osmolarity adjusted to 195 mOsM). Chemical reagents (glycine and picrotoxin) were purchased from Sigma-Aldrich.

### Full-length GlyR cloning and transfection

The zebrafish GlyRα1 shares 92% amino acid similarity with the human GlyRα1. Codon-optimized zebrafish GlyRα1 (NCBI Reference Sequence: NP_571477) was purchased from GenScript. The sequence included the full-length GlyRα1 gene (referred to in the text as GlyR) followed by a thrombin sequence (LVPRGS) and a C-terminal octa-His tag. The gene was subcloned into the pFastBac1 vector for expression in s*podoptera frugiperda* (*Sf9*) cells^60^. The *Sf9* cells were cultured in Sf-900™ II SFM medium (Gibco®) without antibiotics and incubated at 28 °C without CO_2_ exchange. Transfection of sub-confluent cells was carried out with recombinant GlyR bacmid DNA using Cellfectin II transfection reagent (Invitrogen) according to manufacturer’s instructions. Cell culture supernatants were collected and centrifuged at 5 days post-transfection at 1,000g for 5 min to remove cell debris to obtain progeny 1 (P1) recombinant baculovirus. *Sf9* cells were infected with P1 virus stock to produce P2 virus which were then used for subsequent infection to produce P3 viruses and so on till the P6 generation. The P6 virus was used for recombinant protein production based on expression levels and eventually the purified sample quality.

### GlyR expression, purification and nanodisc reconstitution

Approximately 2 × 10^6^ per ml *Sf9* cells were infected with P6 recombinant viruses. After 48 h post-infection, the cells were harvested and centrifuged at 8,000g for 20 min at 4 °C to separate the supernatant from the cell pellet. The cell pellet was resuspended in dilution buffer (20 mM Tris-HCl, pH 7.5, 36.5 mM sucrose) supplemented with 1% protease inhibitor cocktail (Sigma-Aldrich). Cells were disrupted by sonication on ice and non-lysed cells were removed by centrifugation at 3,000g for 15 min. The supernatant was subjected to ultracentrifugation at 167,000g for 1 h to separate the membrane fraction. The membrane pellet was solubilized with 15 mM *n*-dodecyl-β-D-maltopyranoside (DDM, Anatrace) in a buffer containing 200mM NaCl and 20mM HEPES, pH 8.0 (buffer A), supplemental with 0.05 mg/ml soybean polar extract (asolectin, Avanti Polar Lipids) and 0.05% cholesterol hemisuccinate (CHS, Avanti Polar Lipids) for 2 hours at 4°C. Non-solubilized material was removed by ultracentrifugation (167,000g for 25 min). The supernatant containing the solubilized protein was incubated with TALON resin pre-equilibrated with buffer A, 1 mM DDM, 0.05 mg/ml asolectin, and 0.05% CHS for 2 h at 4 °C. The beads were then washed with 10 column volumes of buffer A, 1mM DDM, 0.05 mg/ml asolectin, 0.05% CHS, and 35 mM imidazole. The GlyR protein was eluted with buffer A, 1 mM DDM, 0.05 mg/ml asolectin, 0.05% CHS, and 250 mM imidazole. Eluted protein was concentrated and applied to a Superose 6 column (GE healthcare) equilibrated with buffer A and 1 mM *DDM*. Fractions containing the GlyR pentamers were collected and concentrated to 0.5 mg/ml using 50-kDa MWCO Millipore filters (Amicon). The nanodisc reconstitution was carried out using previously published protocols^61^. Membrane scaffold protein (MSP1E3D1) was expressed and purified as previously described with some modifications^62,63^. The MSP1E3D1 gene in pET 28a (a gift from Stephen Sligar: Addgene plasmid # 20066)^64^. Briefly, the soyabean polar lipid extract (Avanti Lipids) was dried in a stream of nitrogen and equilibrated in buffer A with protein, DDM, and MSP1E3D1 such that final molar ratio of Protein: MSP: lipid: DDM was 1:3:120:5. The mixture was incubated for 1 h with gentle rotation. After 1 hr incubation, Bio-bead SM-2 (Bio-Rad Laboratories) were added and mixture was subjected to gentle rotation for 12 hrs at 4° C. The reconstituted protein was applied to a Superose 6 column (GE healthcare) equilibrated with buffer A. Fractions containing the protein nanodiscs were collected and concentrated to 0.1 mg/ml using 50-kDa MWCO Millipore filters (Amicon) for cryo-EM studies.

### Preparation of sample cryo-EM imaging and parameters for data acquisition

GlyR nanodiscs sample from gel filtration (∼0.1mg/ml) was filtered and used for cryo-EM. For the Apo condition, the sample was used as such without any ligand (GlyR-Apo). For GlyR-Gly and GlyR-Gly/PTX, the samples were incubated for 1 hour with 5 mM glycine and a combination of 5 mM glycine and 3mM picrotoxin (PTX), respectively. The sample was blotted thrice with 3.5 μl sample each time onto glow-discharged Cu 300 mesh Quantifoil 1.2/1.3 grids (Quantifoil Micro Tools) and the grids were plunge frozen into liquid ethane using a Vitrobot (FEI). The grids were imaged using a 300 kV FEI Titan Krios microscope equipped with a Gatan K2-Summit direct electron detector camera (GlyR-Apo and GlyR-Gly/PTX datasets) or Gatan K3 direct electron detector camera for GlyR-Gly dataset. The parameters for data acquisition for the different conditions are as follows. GlyR-Apo: Movies containing 40 frames were collected at 130,000× magnification (set on microscope) in super-resolution mode with a physical pixel size of 1.064 Å/pixel, dose per frame 1.30 e^-^/Å^2^. Defocus values of the images ranged from −1.5 to −2.5 µm (input range setting for data collection) as per the automated imaging software EPU. GlyR-Gly/PTX: 40 frames/movie were collected at 81,000× magnification (set on microscope) in counting mode with a physical pixel size of 1.06 Å/pixel, dose per frame 1.45 e^-^/Å^2^. Defocus values ranged from −1.5 to −2.5 GlyR-Gly: 40 frames/movie were collected at 81,000× magnification (set on microscope) in super-resolution mode with a physical pixel size of 1.08 Å/pixel, dose per frame 1.25 e^-^/Å^2^. Defocus values ranged from −1.5 to −2.5 µm.

### Image processing

MotionCor2^65^ (v 1.2.3) with a B-factor of 300 pixels^2^ was used to correct beam-induced motion. Super-resolution images (for GlyR-Apo and GlyR-Gly datasets) were binned (2×2) in Fourier space, making a final pixel size of 1.064 Å and 1.08 Å respectively. Initially, a small subset of micrographs were CTF-corrected using CTFFIND4^66^ and processed in cisTEM^67^ to generate 2D-template for autopicking. All subsequent data processing was conducted in RELION 3.0^68^. The defocus values of the motion-corrected micrographs were estimated using Gctf software^69^.

For GlyR-Apo dataset: 2, 042 micrographs of a total of 2, 389 were manually sorted. ∼444, 921 auto-picked particles from 2, 042 micrographs using templates generated in cisTEM and were subjected to 2D classification to remove suboptimal particles. 61, 000 good particles were selected and subjected to initial round of 3D auto-refinement using 30 Å low-pass filtered map of related human synaptic α1-β3-γ2 GABAA receptor (EMD-4411) from which Megabody38 was manually erased in chimera. Multiple rounds of 2D classification and 3D classification without image alignment were done to remove low resolution and broken particles. All the particles corresponding to bottom views of receptor were removed from subsequent processing to prevent misalignment of the particles in 3D auto-refinement. Nanodisc belt was subtracted to improve the angular accuracy of alignment and remove sub-optimal particles. A set of 22, 643 particles resulted in 3.7 Å map which was used for training parameters for Bayesian polishing^68,70^. Optimized parameters (s_vel 1.029 --s_div 9585 --s_acc 2.805) were then used to polish these particles and the polished particles were then subjected to multiple rounds of 3D auto-refinement and 3D classification without image alignment. Few rounds of CTF refinement were performed before the final round of 3D auto-refinement. A final subset 19, 653 particles used for auto-refinement, with soft mask accounting for density of nanodisc, resulted in 3.39 Å map which was post-processed resulting in final resolution of 3.36 Å (Fourier shell coefficient -FSC = 0.143 criterion). The B-factor estimation and map sharpening were performed in the post-processing step. Local resolutions were estimated using the RESMAP software^59^.

For GlyR-Gly dataset, a total of 5344 movies were used. For autopicking in RELION 3.0, the final 2D classes from the GlyR-Apo dataset was used as template and a total of 466,000 were auto-picked. A total of 15,000 particles were subjected to independent 3D auto refinement using GlyR-Apo map loss filtered to 20 Å as a template. Multiple rounds of 2D classification, 3D classification without image alignment was done for datasets. All the particles corresponding to bottom views of receptor were removed from subsequent processing to prevent misalignment of the particles in 3D auto-refinement. The final ∼10,000 particles were used for training parameters for Bayesian polishing. Few rounds of CTF refinement and classification were performed before the final round of 3D autorefinement. This resulted in a final map containing 8,255 particles at 3.55 Å after post-processing. The B-factor estimation and map sharpening were performed in the post-processing step. Local resolutions were estimated using the RESMAP software^59^.

For GlyR-Gly-PTX: a total of 2, 280 movies, collected on counting mode, were used from two imaging sessions. The grids for these imaging sessions were from the same batch, collected on the same microscope using identical parameters, For auto-picking in RELION3.0, the final 2D classes from the GlyR-Apo dataset was used as template and a total of 400,000 particles were picked. A total of 8, 288 particles from dataset 1 (1, 470 movies) and 11, 331 particles from dataset 2 (810 movies) were subjected to independent 3D autorefinement using GlyR-Apo map loss filtered to 20 Å as a template. Multiple round of 2D classification and 3D classification without image alignment was used to get rid of low resolution particles. All the particles corresponding to bottom views of receptor were removed from subsequent processing to prevent misalignment of the particles in 3D auto-refinement. The two datasets were individually used for Bayesian polishing and the final polished datasets were merged in RELION 3.0.6. Few rounds of CTF refinement was performed before the final round of 3D autorefinement. Merged dataset with 12, 503 particles was used for 3D autofinement and further classification and resulted in a final map containing 10, 375 particles at 3.65 Å after post-processing. The B-factor estimation and map sharpening were performed in the post-processing step. Local resolutions were estimated using the RESMAP software^59^.

### GlyR model building

The 3D cryo-EM maps for the GlyR-Apo, GlyR-Gly, and GlyR-Gly-PTX data sets used for model-building contained density for the entire ECD, TMD and a small region of the ICD. The final refined models comprised of residues Pro31–Phe341 and Lys394–Gln444. The missing region (342–393) is of the unstructured ICD. The previously solved structure of GlyR-strychnine (PDB ID: 3JAD) was used as a starting model for the GlyR-Apo data set. The residues were renumbered as per the sequence available in UniProt database (O93430). For model building of GlyR-Gly and GlyR-Gly-PTX, the GlyR-Apo structure was used as a template. The cryo-EM map was first converted to the .mtz format using CCP4i software^71^ with mapmask and sfall tools and then used for manual model building in COOT^72^. After initial model building, the three states were refined against their corresponding EM-derived maps using the phenix.real_space_refinement tool from the PHENIX software package^73^, using rigid body, local grid, NCS, and gradient minimization. The individual models were then subjected to additional rounds of manual model fitting and refinement. The refinement statistics, the final model to map cross-correlation evaluated using phenix module mtriage^74^, and the stereochemical properties of the models as evaluated by Molprobity^75^ are detailed in the Supplemental Table 1. Protein surface area and interfaces were analyzed using the PDBePISA server (http://www.ebi.ac.uk/pdbe/pisa/). The pore profile was calculated using the HOLE program^56^. The cavities were analyzed using Fpocket algorithm^58^. Figures were prepared using PyMOL v.2.0.4 (Schrödinger, LLC).

**Supplemental Table 1.**

|  | **GlyR-Apo**<br>**(6UBS, 20714)** | **GlyR-Gly**<br>**(6UBT, 20715)** | **GlyR-Gly-PTX**<br>**(6UD3, 20731)** |
| --- | --- | --- | --- |
| **Data Collection and processing** |  |  |  |
| Microscope and location | FEI Titan Krios,<br>Frederick<br>National<br>Laboratory for<br>Cancer Research | FEI Titan Krios,<br>Frederick National<br>Laboratory for<br>Cancer Research | FEI Titan Krios,<br>Stanford-SLAC<br>Cryo-EM Center |
| Magnification | 130000 | 81000 | 81000 |
| Voltage | 300 | 300 | 300 |
| Detector | super-resolution<br>mode | super-resolution<br>mode | Counting mode |
| Physical pixel size | 1.06 Å/pixel | 1.08 Å/pixel | 1.06 Å/pixel |
| Defocus range (µM) | -1.5 to -2.5 | -1.5 to -2.5 | -1.5 to -2.5 |
| Number of images | 2389 | 5344 | 2280 |
| Number of frames/image | 40 | 40 | 40 |
| Initial particle number | ~444,921 | ~466,000 | ~400,000 |
| Final particle number | 19653 | 8255 | 10, 375 |
| Symmetry | C5 | C5 | C5 |
| Resolution (unmasked, Å) | 3.89 | 4.3 | 4.23 |
| Resolution (masked, Å) | 3.33 | 3.55 | 3.63 |
| Map resolution range * | 3-7 | 3-7 | 3-7 |
| Map sharpening B-factor (Å<sup>2</sup>) | -50 | -30 | -20 |
| **Refinement** |  |  |  |
| Initial model used (PDB code) | 3JAD | NA | NA |
| Composition |  |  |  |
| Protein residues | 1810 | 1810 | 1810 |
| Non Hydrogen atoms | 15465 | 14965 | 14986 |
| Number of atoms Protein Ligands |  |  |  |
| Glycan (NAG) (molecule) | 10 | 10 | 10 |
| Glycan ( BMA) (molecule) | 5 | 5 | 5 |
| PX4 (molecule) | 10 | 0 | 0 |
| PIO (molecule) | 5 | 0 | 0 |
| Glycine (molecule) | 0 | 5 | 5 |
| PTX(RI5) (molecule) | 0 | 0 | 1 |
| Bonds (RMSD) |  |  |  |
| Length (Å) (# > 4σ) | 0.003 (0) | 0.003 (0) | 0.004 (0) |
| Angles (°) (# > 4σ) | 0.743 (20) | 0.747 (5) | 1.043 (11) |
| Ramachandran plot (%) |  |  |  |
| Outliers | 0 | 0 | 0 |
| Allowed | 4.47 | 4.19 | 2.51 |
| Favored | 95.53 | 95.81 | 97.49 |
| Rotamer outliers (%) | 0.31 | 0.31 | 0.12 |
| Molprobity score | 1.6 | 1.61 | 1.35 |
| Molprobity clashscore | 5.31 | 5.9 | 4.79 |

\* Local resolution range

### Molecular Dynamics Simulations

Molecular structures of the GlyR in various conformational states were separately embedded within phospholipid (POPC, 1-palmitoyl-2-oleoyl-*sn*-glycero-3-phosphocholine) bilayer membranes that were solvated on either side to form 13 x 13 x 17 nm^3^ simulation cells. Each protein-membrane system was assembled and equilibrated via a multiscale protocol^76^. Simulations were performed with GROMACS 2018^77^, using the TIP4P/2005 water model^78^ and the OPLS all-atom protein force field with united-atom lipids^79^. The integration time-step was 2 fs. Bonds were constrained through the LINCS algorithm^80^. A Verlet cut-off scheme was applied, and long-range electrostatic interactions were calculated using the Particle Mesh Ewald method^81^. Temperature and pressure were maintained at 37 °C and 1 bar during simulations, using the velocity-rescaling thermostat^82^ in combination with a semi-isotropic Parrinello and Rahman barostat^83^, with coupling constants of τ_T_ = 0.1 ps and τ_P_ = 1 ps, respectively.

Pore water free energy profiles were computed for alternative conformations of the protein using the Channel Annotation Package^37^, in each case based on three replicates of 30 ns equilibrium simulations at physiological salt (150 mM NaCl) concentration. To preserve the conformational state of each cryo-EM structure (while permitting rotameric flexibility in amino acid side chains) during simulations, harmonic restraints at a force constant of 1000 kJ mol^-1^ nm^-2^ were placed on protein backbone atoms. Simulation trajectories were analyzed at 500 ps intervals, with a bandwidth of 0.14 nm applied for water density estimation.

Chloride conduction was measured in five replicates of 200 ns simulations for each (backbone-restrained) receptor structure, at 500 mM NaCl concentration and in the presence of a 200 mV transmembrane potential difference, with negative potential on the cytoplasmic side. This was applied by imposing an external, uniform electric field in the membrane normal direction. Conductance values were calculated from the number of permeation events and averaged between replicates.

A separate set of equilibrium simulations (at 150 mM NaCl, in the absence of a transmembrane potential difference) were used to monitor the behavior of each protein conformation by sequential relaxation followed by removal of positional restraints. During an initial 10-ns simulation, harmonic restraints (again with force constant 1000 kJ mol^-1^ nm^-2^) were placed on all non-hydrogen atoms of the protein. Restraints on side chain atoms were then released, and the protein subjected to a further 20 ns of simulation. In the next 20 ns, only the α-carbon atom of each residue remained under the positional restraining force. The final state of the system was subsequently simulated for 200 ns, unrestrained.

### Data Availability Accession Numbers

The atomic coordinates and cryo-EM maps for the GlyR-Apo, GlyR-Gly, and GlyR-Gly/PTX structures have been deposited in the Protein Data Bank and Electron Microscopy Data Bank with accession codes PDB-6UBS (EMD-20714), PDB-6UBT (EMD-20715) and PDB-6UD3 (EMD-20731), respectively. All relevant data are available from the corresponding author upon reasonable request.

## Acknowledgements

We acknowledge the use of instruments at the National Cryo-Electron Microscopy Facility at the NCI and Stanford-SLAC Cryo-Electron Microscopy Facility. A special thanks to Prof. Wah Chiu for providing us imaging time and supervising the data collection (supported by National Institutes of Health grants: P41GM103832 and S10OD021600). We are grateful to the Cryo-Electron Microscopy Core at the CWRU School of Medicine and Dr. Kunpeng Li for the access to the sample preparation and Cryo-EM instrumentation. We thank Dr. Walter F. Boron for kindly providing us *Xenopus* oocytes and for unrestricted access of the oocyte rig. We are deeply appreciative of the support provided by Dr. Fraser Moss and Mr. Brian Zeise with the oocyte rig. We are very grateful to the members of the Chakrapani lab for critical reading and comments on the manuscript. This research was, in part, supported by the National Cancer Institute’s National Cryo-EM Facility at the Frederick National Laboratory for Cancer Research. This work was supported by the National Institutes of Health grants R01GM108921, R01GM131216, and Cryo-EM supplements: 3R01GM108921-03S1, R01GM108921-5S1, 3R01GM131216-1S1 to S.C and the AHA postdoctoral Fellowship to S.B (17POST33671152).

## Author Contributions

A.K and S.C conceived the project and designed experimental procedures. A.K purified the protein, and with assistance from S.B, optimized the cryo-EM sample preparation and performed grid screening. A.K carried out cryo-EM data analysis, model building, and refinement with inputs from S.B. M.L.M collected the cryo-EM data for GlyR-Gly/PTX. A.K and Y.G performed two-electrode voltage-clamp recordings. S.R performed the MD simulations, under the supervision of M.S. S.C supervised the execution of the experiments, data analysis, and interpretation. A.K and S.C drafted the manuscript with contributions from S.B, S.R, Y.G, and M.S. All authors reviewed the final manuscript.

## Supplemental Information

**Supplemental Figure 1.**
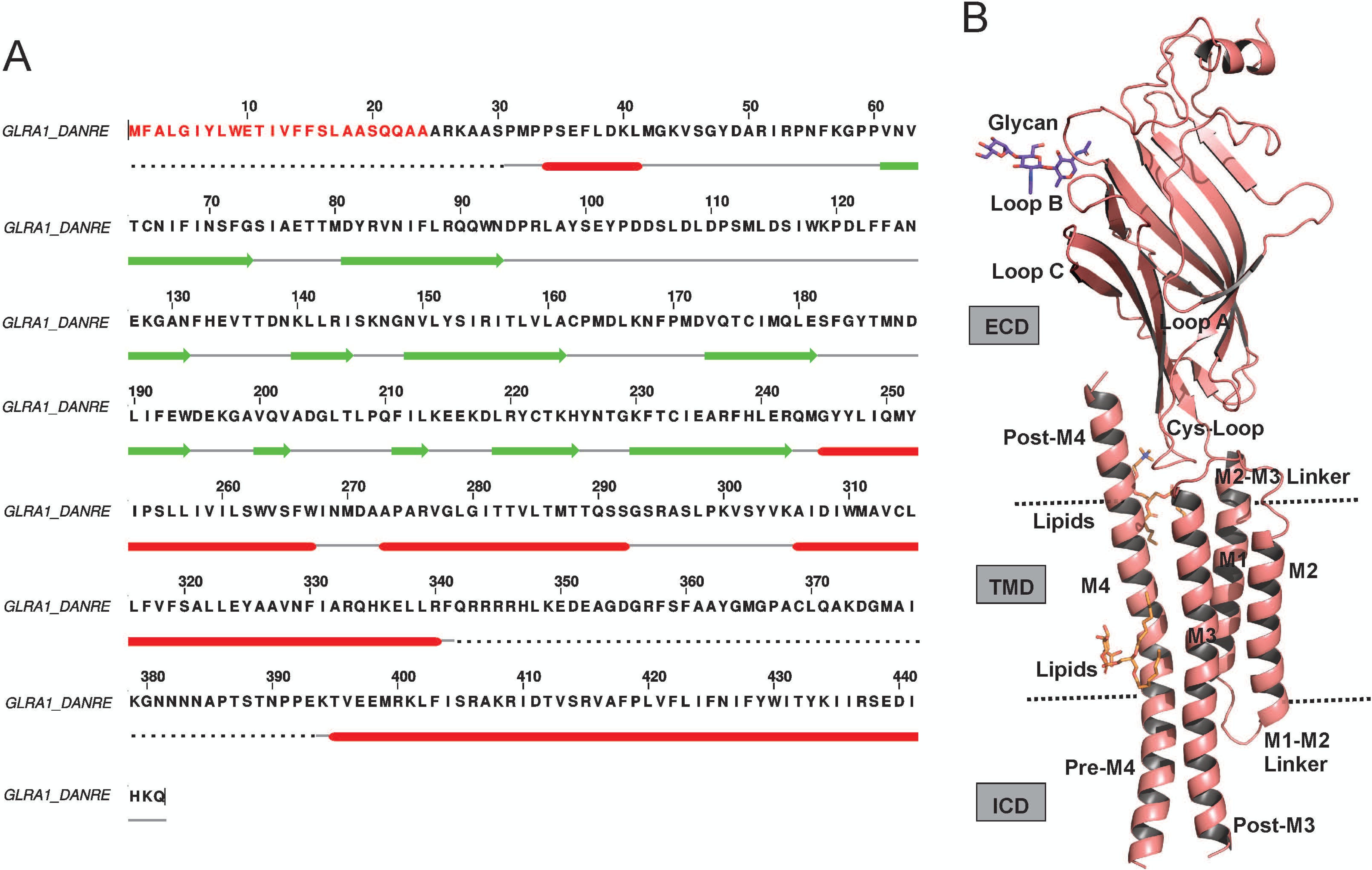
Sequence and topology of GlyR. A) Sequence of zebrafish GlyRα1 used in the cryo-EM study and electrophysiological analysis. Secondary structural elements are indicated above the sequence. B) A single-subunit of GlyR-Apo, viewed from a plane parallel to the membrane, with secondary structure elements labeled. The glycans are shown as magenta sticks and lipids are shown in orange sticks. The putative membrane limits are marked by dotted lines.

**Supplemental Figure 2.**
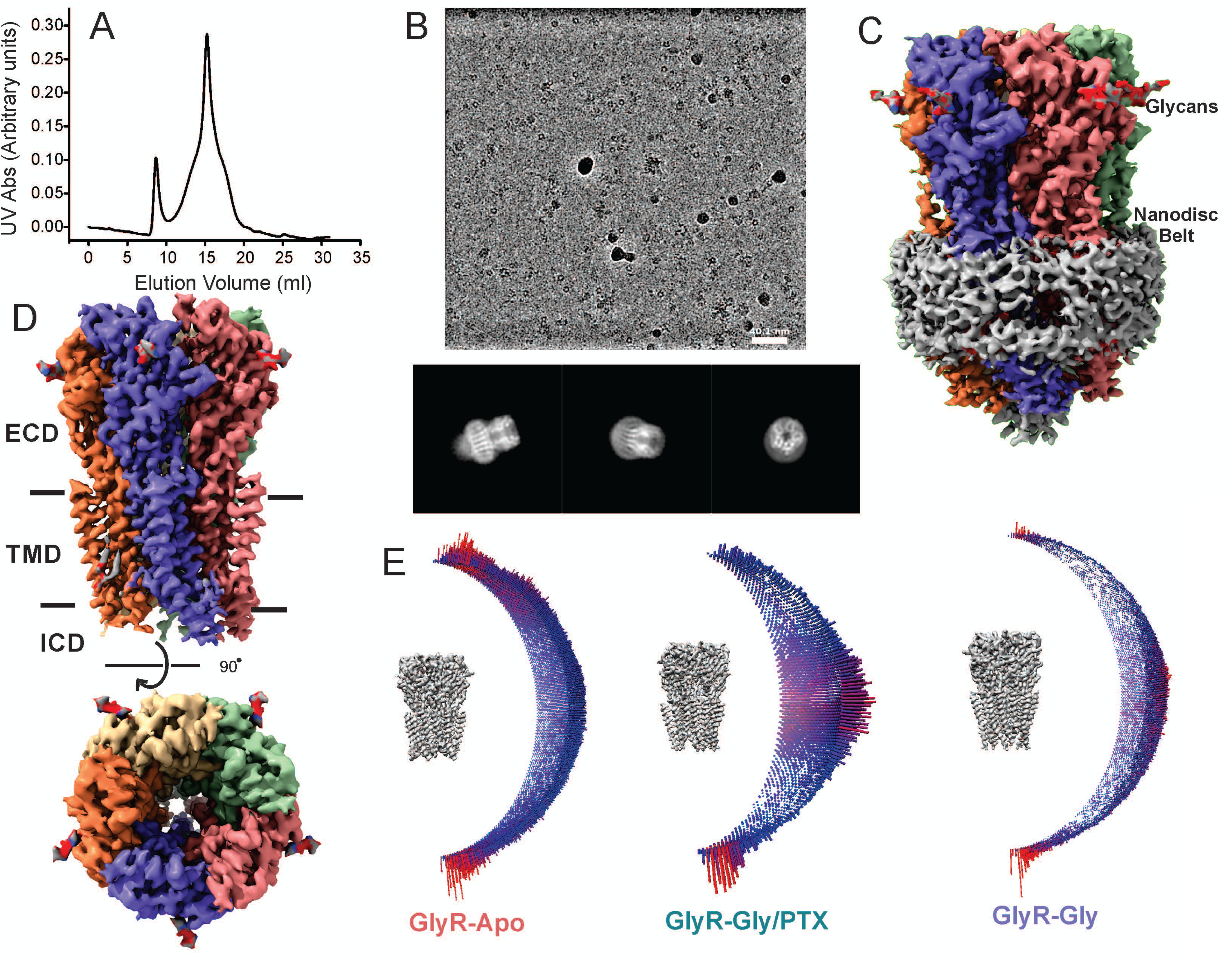
Biochemical and structural characterization of GlyR. A) Gel-filtration profile of GlyR expressed and affinity-purified from *Sf9* cells on a Superose 6 Increase 10/300 GL column (GE Healthcare). The main peak corresponds to the GlyR pentamers. B) A representative cryo-EM micrograph of nanodisc-reconstituted GlyR-Apo sample in vitreous ice (*top*) and selected 2D classes showing various orientations (*bottom*). **C)** Cryo-EM 3D reconstruction of GlyR-Apo. Each subunit is individually colored for clarity and the glycans are show in red. The density corresponding to the nanodisc belt is colored gray. D) Side and top views of the 3D reconstruction showing GlyR-Apo after subtracting the nanodisc belt. The individual domains are marked. E) Angular distribution of particle projections for the final reconstruction used for model building. The map of the GlyR-Apo, GlyR-Gly and GlyR-Gly/PTX complex is shown in gray. Nanodisc belts have been removed for clarity. Length of each cylinder corresponds to the number of particles at a specific Euler angle.

**Supplemental Figure 3.**
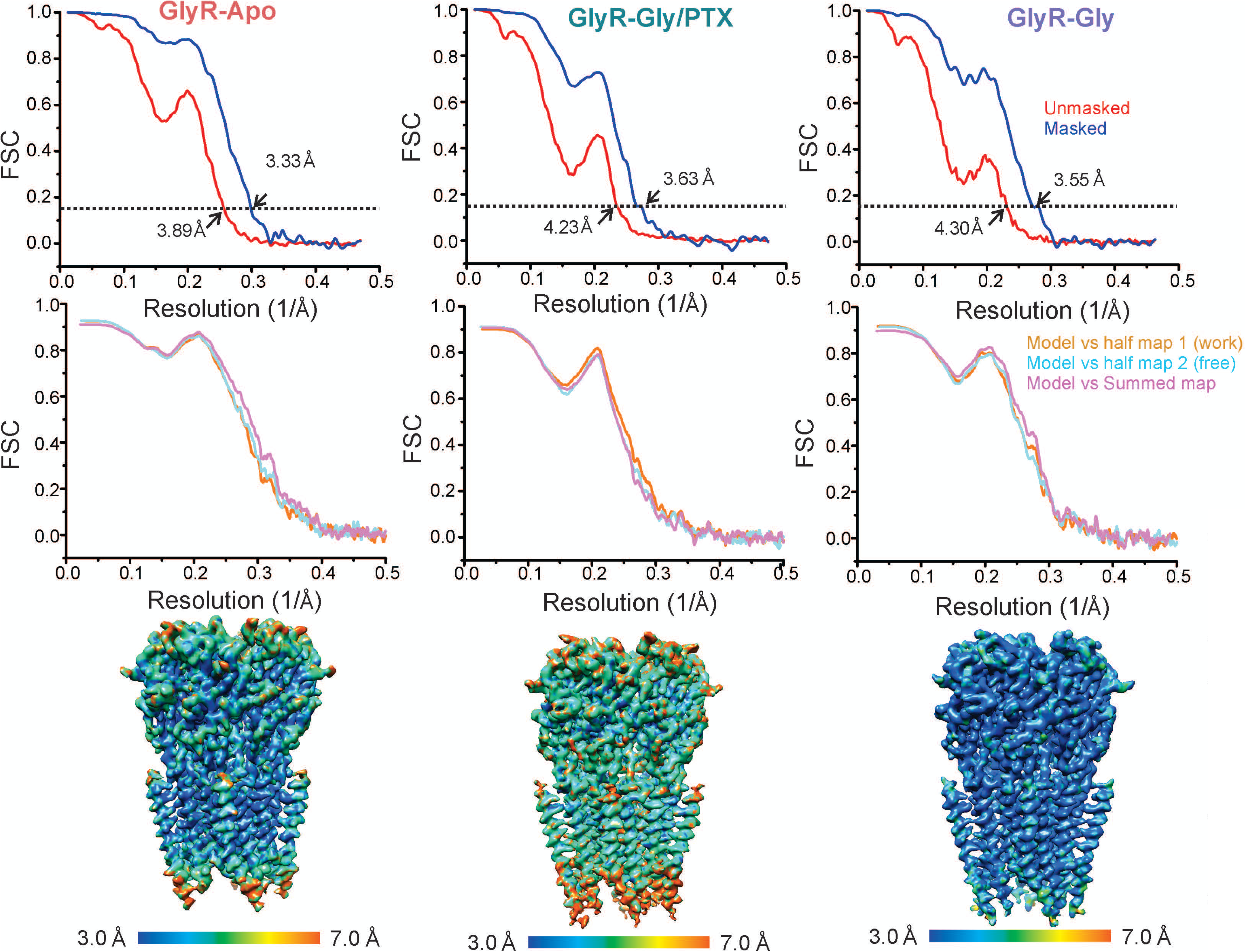
Resolution estimation and model validation. Fourier shell correlation (FSC) curves before (red) and after post-processing (blue) using gold-standard refinement in RELION 3.0 *(top)*. The dashed line represents an FSC of 0.143. For cross validation of model refinement, FSC curves of the refined model versus summed map (full dataset), refined model versus half map 1 (used during refinement), and refined model versus half map 2 (not used during refinement) *(middle)*. Side views of the 3D reconstructions colored-coded by the local resolution determined using ResMap program algorithm v1.1.5 ^59^ *(bottom)*.

**Supplemental Figure 4.**
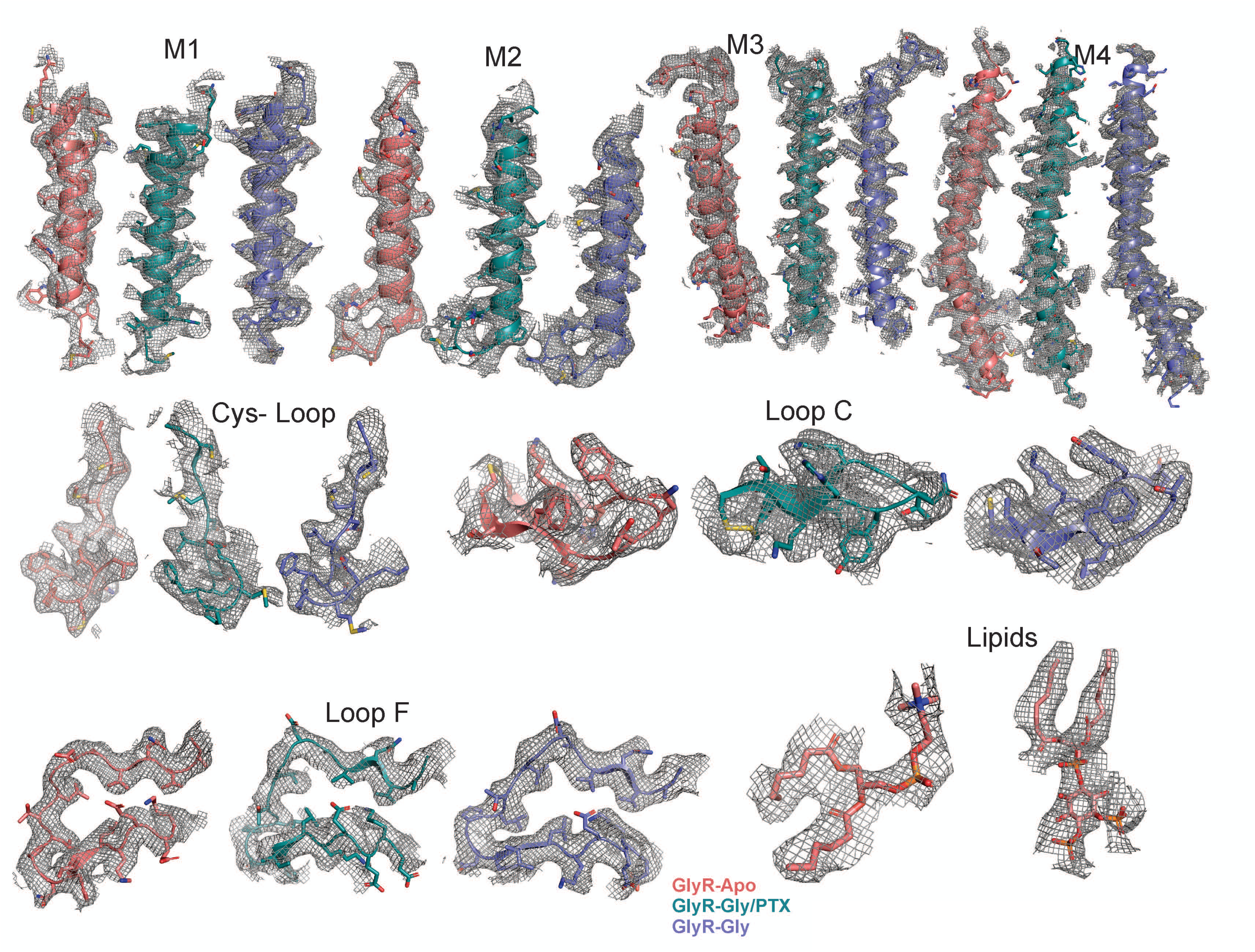
Map correlation of GlyR structures. Validation of various regions within each of the domains of the model (shown as cartoon with stick representation for the residues) and corresponding density map (mesh) are shown here. The contour levels for depicted regions in GlyR-Apo: M1 (6σ), M2 (6σ), M3 (6σ), M4 (5σ), Cys loop (7σ), Loop C (8σ), Loop F (8σ), Lipids (5σ). For GlyR-Gly, the contour levels: M1 (6σ), M2 (5σ), M3 (4.5σ), M4 (6σ), Cys loop (7σ), Loop C (8σ), Loop F (8σ). For GlyR-Gly-PTX, the contour levels: M1 (7σ), M2 (5σ), M3 (5σ), M4 (6σ), Cys loop (6σ), Loop C (8σ), Loop F (9σ).

**Supplemental figure 5.**
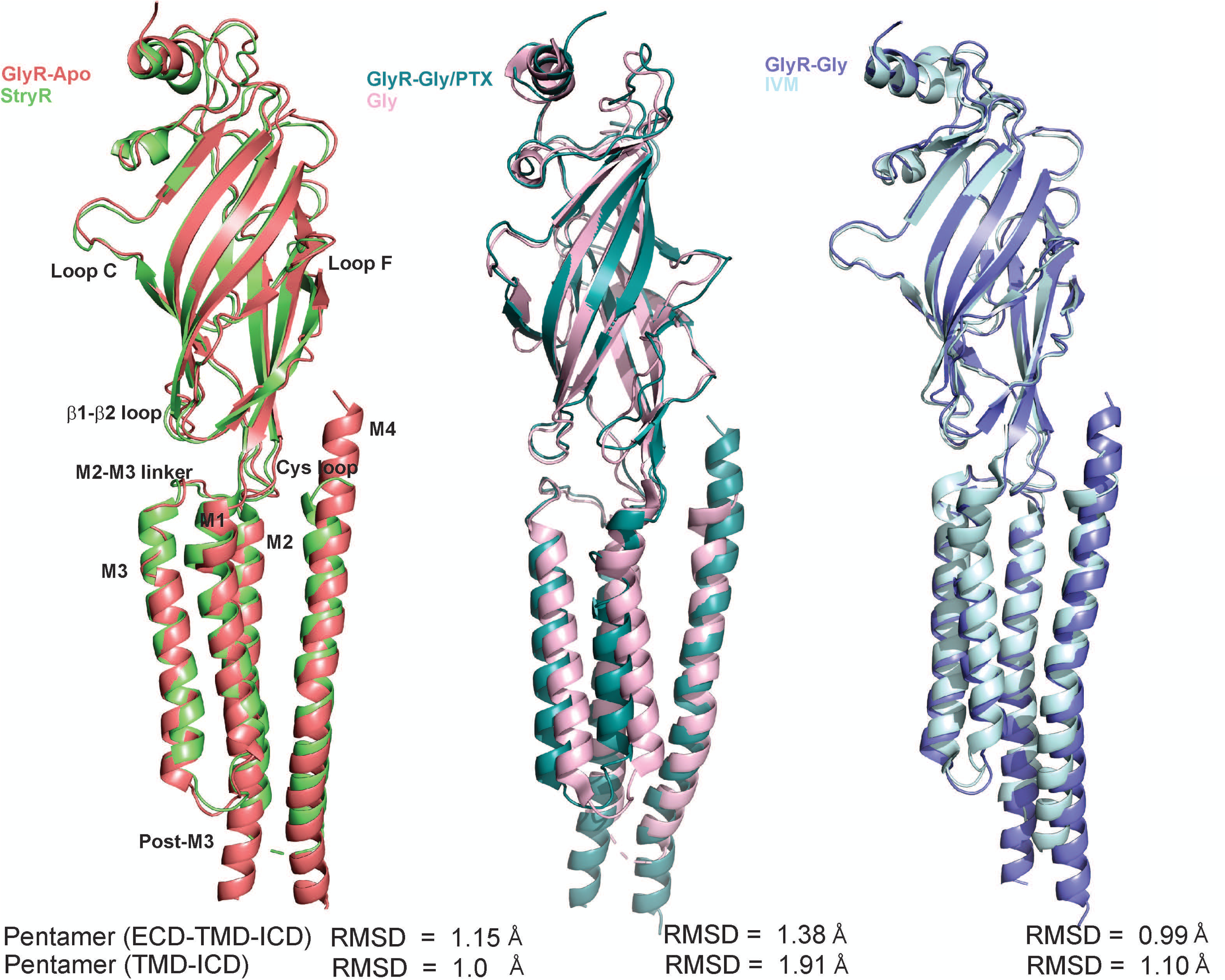
Comparison with previous GlyR cryo-EM structures. Superposition of the single subunits of GlyR-Apo with GlyR strychnine-bound state (PDB_ID:3JAD), GlyR-Gly-PTX with GlyR Gly-bound open state (PDB_ID:3JAE) and GlyR-Gly with GlyR ivermectin-bound desensitized state (PDB_ID:3JAF)^6^. The RMSD (alignment of the pentamers) were calculated to be 1.149Å, 1.381Å, and 0.993 Å respectively.

**Supplemental Figure 6.**
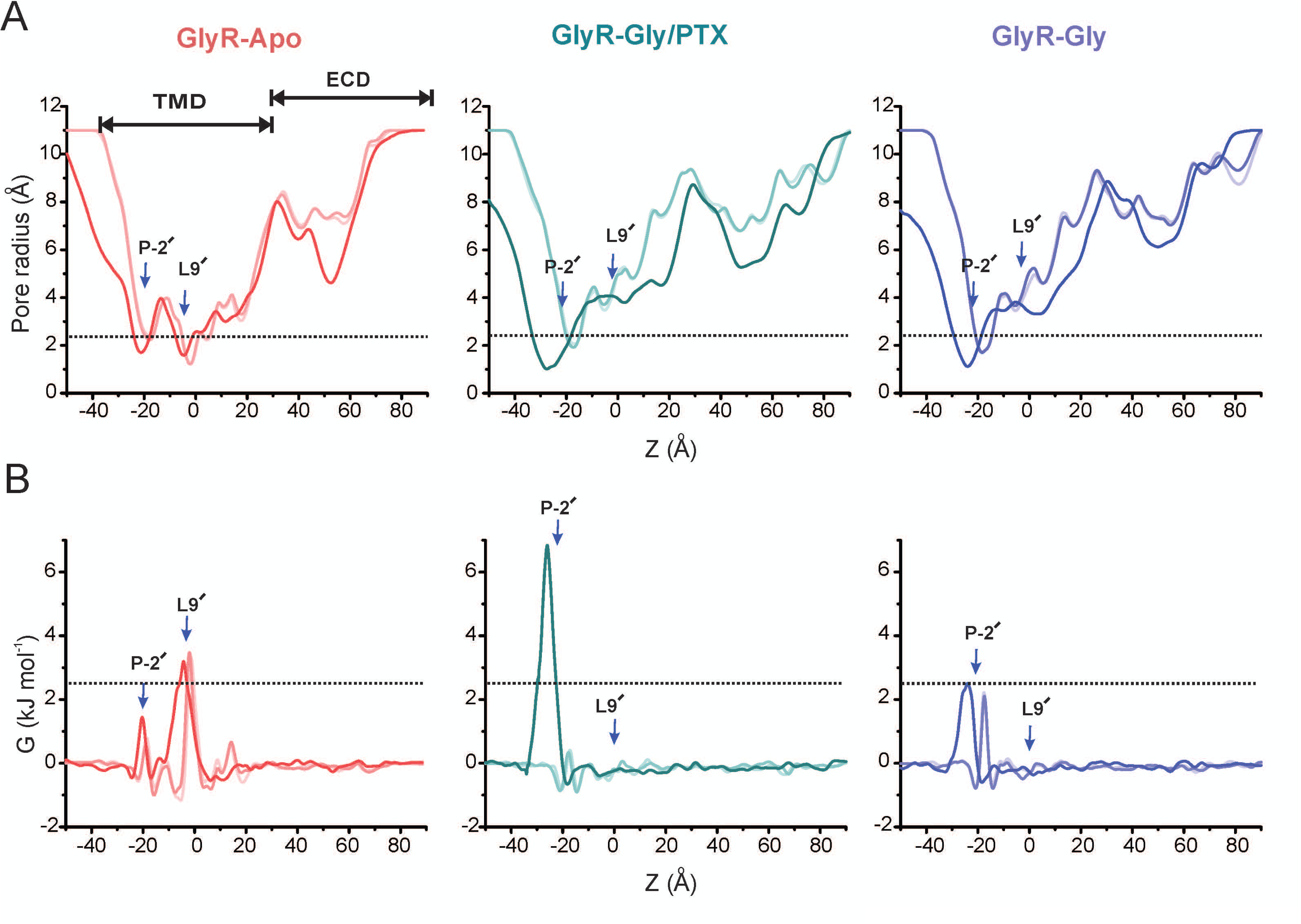
Pore profiles during and after sequential relaxation and removal of protein restraints. A) Radius and B) water free energy profiles determined during a series of simulations in which positional restraints were released successively from the protein side chains (lightest colored lines), from all non-Cα atoms (darker lines), and then from all atoms (i.e. with the protein entirely unrestrained; darkest lines). Each profile was averaged over the final 20 ns of simulation trajectory during that stage.

**Supplemental Figure 7.**
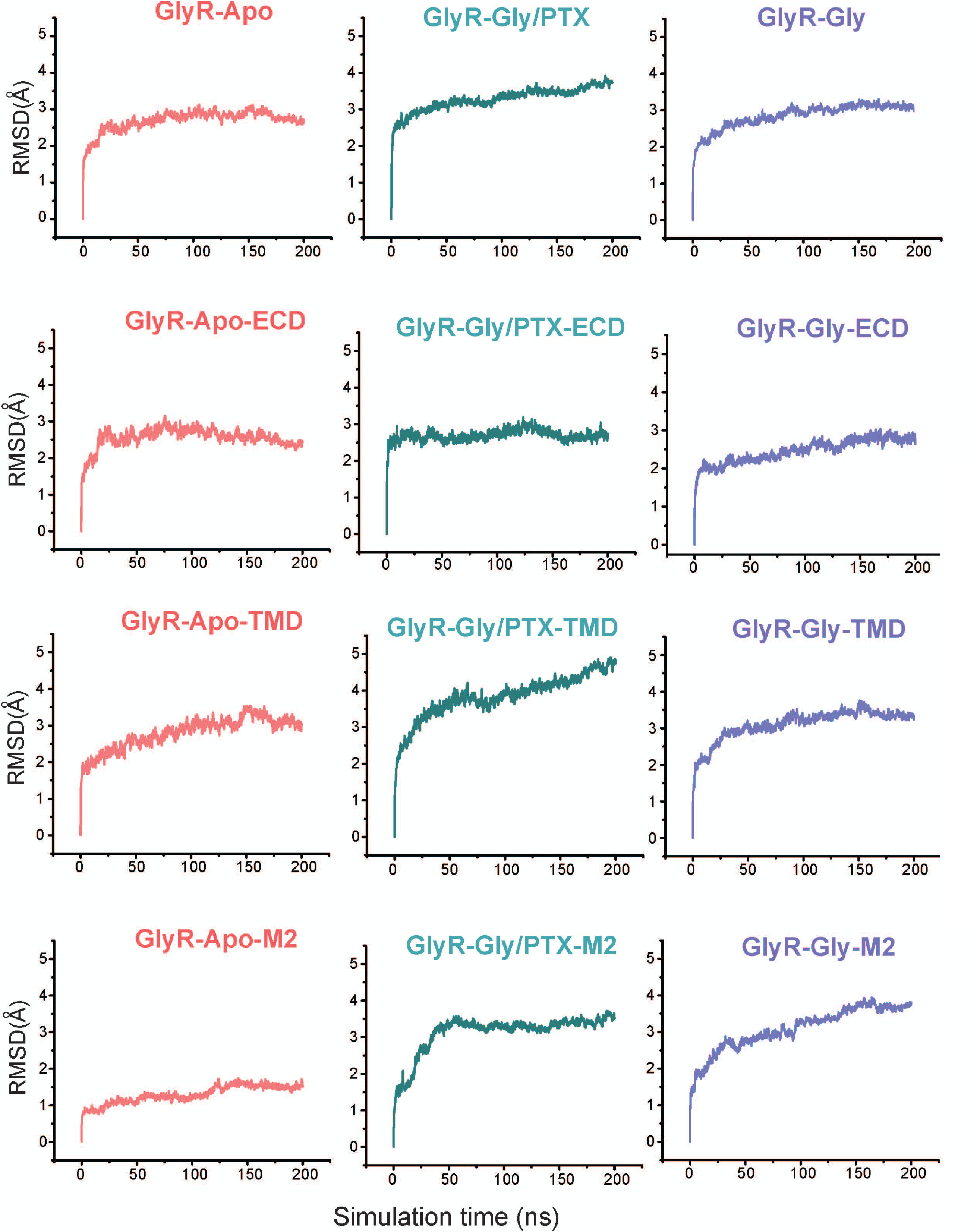
Root-mean-square deviations of protein Cα atoms during unrestrained simulations. Measurements begin upon removal of all restraints on the protein, sampling every 100 ps. The first row showing RMSD for the entire structure, the second row for the ECD, the third for the TMD, and the bottom row for that of M2.

**Supplemental Movie 1. Conformational changes underlying GlyR gating.** A morph of the GlyR-Apo, GlyR-Gly/PTX, and GlyR-Gly structures to highlight the global conformational changes that ensues upon glycine binding in all three domains of the channel.

**Supplemental Movie 2. Trajectory of a single chloride ion (purple sphere) traveling through the GlyR-Gly/PTX structure, in the presence of a transmembrane potential difference of 200 mV.** The duration of the movie corresponds to ∼30 ns of a 200 ns simulation. Water molecules within 4Å of the ion at a given instant are shown as red-and-white sticks; all other water and ions present in the system are omitted for clarity. Two subunits of the protein (semi-transparent surface) are shown in cartoon representation. Membrane lipid head-groups are colored orange.

